# High-throughput nanopore DNA sequencing of large insert fosmid clones directly from bacterial colonies

**DOI:** 10.1101/2024.02.05.578990

**Authors:** Léa Chuzel, Amit Sinha, Caileigh V. Cunningham, Christopher H. Taron

**Affiliations:** New England Biolabs, 240 County Road, Ipswich, MA 01938, USA; Gloucester Marine Genomics Institute, 417 Main Street, Gloucester, MA 01930

## Abstract

Fosmids and cosmids are vectors frequently used in functional metagenomic studies. With a large insert capacity (around 30 kb) they can encode dozens of cloned genes or in some cases, entire biochemical pathways. Fosmids with cloned inserts can be transferred to heterologous hosts and propagated to enable screening for new enzymes and metabolites. After screening, fosmids from clones with an activity of interest must be *de novo* sequenced, a critical step towards identification of the gene(s) of interest. In this work, we present a new approach for rapid and high-throughput fosmid sequencing directly from *Escherichia coli* colonies without liquid culturing or fosmid purification. Our sample preparation involves fosmid amplification with phi29 polymerase and then direct nanopore sequencing using the Oxford Nanopore Technologies system. We also present a bioinformatics pipeline termed “phiXXer” that facilitates both *de novo* read assembly and vector trimming to generate a linear sequence of the fosmid insert. Finally, we demonstrate accurate sequencing of 96 fosmids in a single run and validate the method using two fosmid libraries that contain cloned large insert (∼30-40 kb) genomic or metagenomic DNA.

**Importance:** Large-insert clone (fosmids or cosmids) sequencing is challenging and arguably the most limiting step of functional metagenomic screening workflows. Our study establishes a new method for high-throughput nanopore sequencing of fosmid clones directly from lysed *E. coli* cells. It also describes a companion bioinformatic pipeline that enables *de novo* assembly of fosmid DNA insert sequences. The devised method widens the potential of functional metagenomic screening by providing a simple, high-throughput approach to fosmid clone sequencing that dramatically speeds the pace of discovery.

## Introduction

The diversity of microorganisms on earth is estimated to be over one trillion species (1). Microbes have been found in all environments including those with extreme conditions (*e*.*g*., deep sea hydrothermal vents or soda lakes) (2–4). To thrive in each environment, microbes have adapted by producing metabolites and enzymes that are advantageous to their survival in their surroundings. Consequently, Earth’s vast microbiome represents an incredible resource for discovery of new biochemical pathways, enzymes, or metabolites. However, exploring this genetic resource can be challenging as only 1-15% of microorganisms can be cultivated in the laboratory (5, 6). To overcome this challenge, the field of metagenomics uses culture-independent methods whereby microbial DNA is directly isolated from an ecosystem sample (referred to as environmental DNA [eDNA]) (7, 8). In sequencing-based metagenomics, eDNA is directly sequenced, while in functional-metagenomic studies eDNA is cloned in a heterologous host and subsequently screened for expression of desired biochemical activities using high-throughput assays. Cosmids or fosmids (cosmids with inducible copy number) vectors are typically used due to their large insert capacity (about 30-40 kb) that enables many genes to be screened at once (9, 10). Because functional metagenomics detects evidence of gene function, it can be used to assign function to genes previously annotated as ‘hypothetical proteins’ and to establish new protein families in an unbiased manner (11, 12). During screening, clones displaying an activity of interest are identified, after which their fosmids are isolated for sequencing. In large high-throughput screens, this necessitates isolation of fosmids from dozens to hundreds of bacterial clones, a time-consuming, cumbersome, and costly endeavor that slows the pace of discovery. Rapid and high-throughput methods to sequence fosmid clones are needed to overcome this bottleneck.

There are several challenges associated with sequencing fosmid clones. First, fosmid DNA must be isolated from its surrogate host for each clone of interest. Fosmids are typically extracted from liquid bacterial cultures using commercially available fosmid isolation kits. Common plasmid miniprep kits can also be used but are not optimal to obtain high yield and high quality fosmid DNA due to the higher molecular weight of fosmid clones. While these methods are effective, they are cumbersome and costly when applied to the large number of clones high-throughput screens identify. High-throughput fosmid isolation has also been reported using liquid handling robotic platforms with specific automation (13). Once fosmid DNA has been isolated, multiplexing strategies must be employed to tag each fosmid individually to enable sequencing of multiple fosmids at once. Current barcoding workflows are time-consuming and expensive with limited throughput (14). Finally, reads obtained from sequencing must be assembled to obtain the entire fosmid sequence. Due to the metagenomic origin of cloned fosmid inserts, many insert sequences will not map to known DNA from previously sequenced species. Thus, sequencing reads need to be assembled *de novo* in the absence of reference data.

In this study, we describe a new approach for high-throughput sequencing of fosmids on the Oxford Nanopore technology (ONT) platform. ONT is a nanopore-based sequencing technology that can generate long-reads, making it particularly suited for *de novo* DNA sequencing and assembly (15, 16). To overcome the low throughput associated with fosmid DNA isolation from bacterial cultures, our workflow utilizes fosmid amplification directly from lysed bacterial colonies. Our approach exploits the high processivity and strong strand-displacement activities of the phi29 polymerase family (17) to specifically amplify fosmids ahead of barcoding for ONT sequencing. Additionally, a customized bioinformatics pipeline (termed phiXXer, available at https://github.com/aWormGuy/phiXXer) is described that performs automated *de novo* assembly of ONT long-reads from phi29-amplified DNA. This high-throughput workflow significantly simplifies fosmids sequencing and provides a faster route to discovery using functional metagenomics.

## Results

### 1. Amplification of fosmids from colonies using phi29-XT

We sought to develop a high-throughput fosmid sequencing sample preparation method that would circumvent the need to isolate fosmids from bacterial clones (Figure 1). To accomplish this, we amplified fosmids directly from a small volume of crude lysate using phi29 DNA polymerase. This enzyme has high-fidelity and processivity over 70 kb (17). Phi29 is a bacteriophage polymerase that is typically employed in isothermal rolling circle amplification (RCA) methods to amplify circular DNA. In such applications, it is used in combination with random hexamers to permit amplification of any DNA sequence, including whole genomes or circular molecules (18). In our application, we instead used two primers (forward and reverse) specific to the fosmid backbone, to enable fosmid-specific amplification without amplification of host chromosomal DNA.

**Figure 1.**
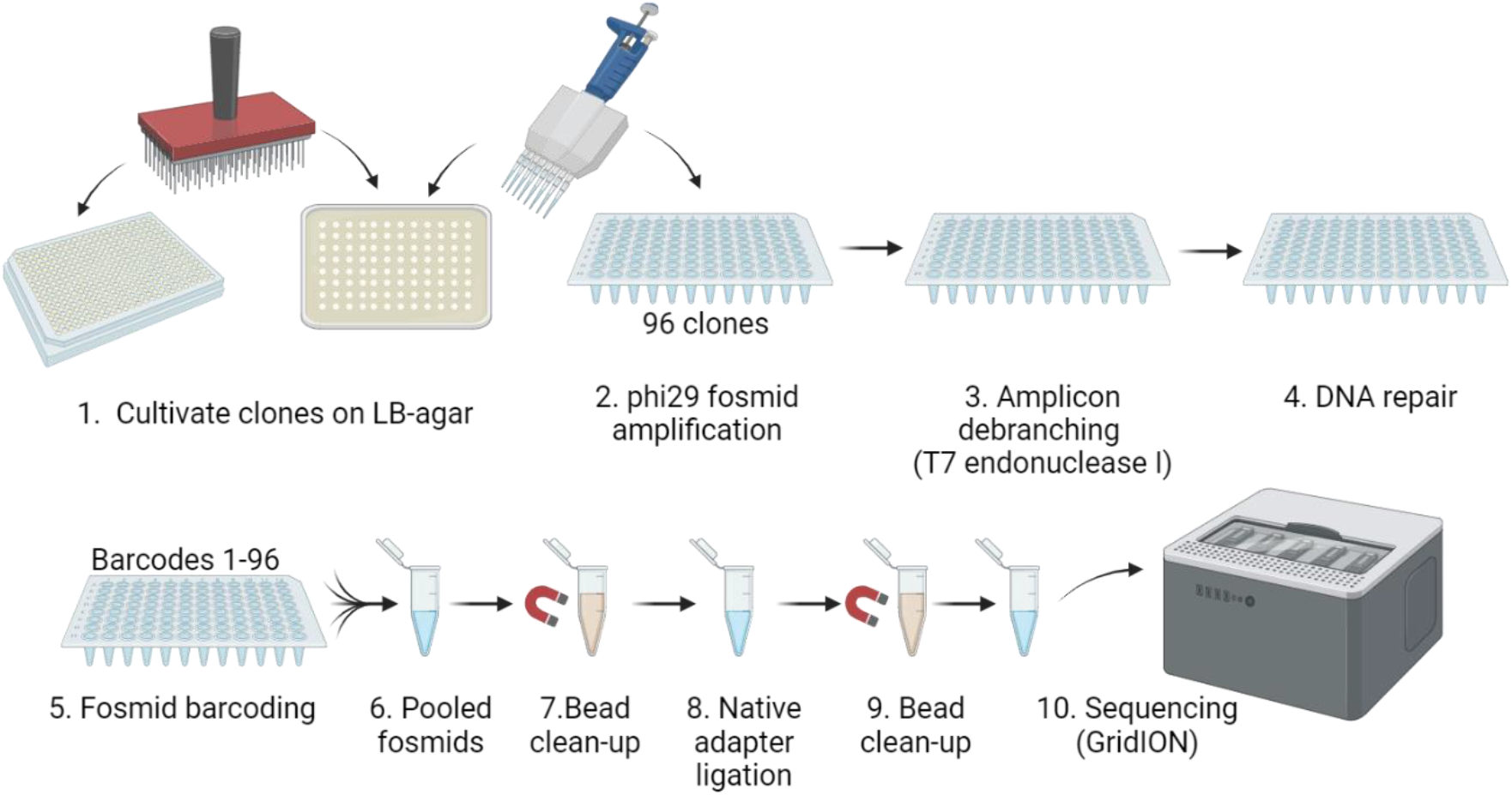
Workflow for high-throughput nanopore sequencing of fosmid clones. The illustrated workflow enables sequencing of 96 fosmids in a single sequencing run using Oxford Nanopore Technologies. Created with BioRender.com.

To illustrate the robustness of isothermal fosmid amplification, fosmids from a *Thermococcus kodakarensis (T. kodakarensis)* genomic library were amplified using phi29-XT DNA polymerase. The *T. kodakarensis* fosmid library was created previously by cloning ∼40 kb fragments of *T. kodakarensis* genomic DNA into pCC1FOS vector and using them to transform *Escherichia coli* EPI300 cells (19). Individual clones were then arrayed in a 384-well format. This fosmid library wa*s* formerly used in various functional screening assays and was originally sequenced and assembled from Illumina sequencing of libraries constructed using the plexWell™ 384 Library Preparation kit (seqWell, Danvers, MA) with fosmid DNA purified from liquid cultures as the input material (19). These sequences serve as a reference to demonstrate the efficacy of our workflow.

In this experiment, 96 arrayed colonies from the *T. kodakarensis* library were grown on LB-agar plates. Cells were introduced into a fresh 96-well PCR plate for cell lysis and 8 h amplification as described in the Materials and Methods. Reaction products were analyzed by restriction enzyme digestion and fragment separation by gel electrophoresis (Figure 2). The distinct banding pattern observed was consistent with digested fosmid DNA. The amplification yield was qualitatively consistent for nearly all fosmids, with only five showing weak or no amplification (fosmids #7, 76, 80, 81 and 90) (Figure 2). Amplification reactions run for a shorter time were also tested (2 h and 4 h) but yielded significantly less DNA (Supplementary figure 1). Therefore, 8 h amplification was considered optimal from the conditions we tested.

**Figure 2.**
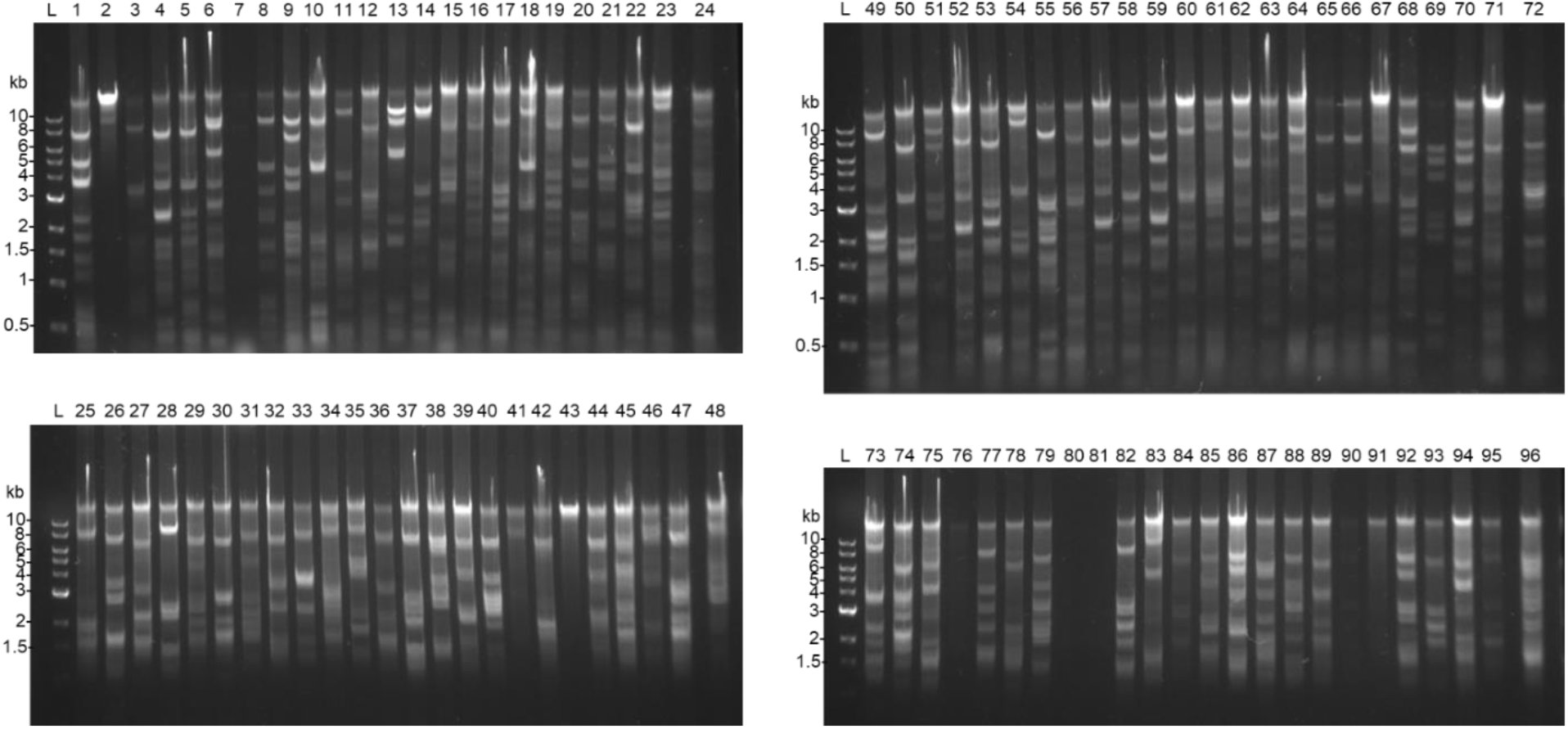
Amplification of 96 fosmids using phi29-XT. A total of 96 clones (lanes 1-96) from a *T. kodakarensis* genomic fosmid library were amplified using phi29-XT. Amplification products were digested with SacI and XhoI and digested products analyzed by separation on a 0.8% agarose gel.

### 2. A nanopore sequencing pipeline for phi29-amplified fosmids

Phi29-amplified fosmids were subjected to nanopore sequencing using ONT. Phi29 polymerase is known to produce complex hyper-branched structures and chimeras in amplified DNA (20). These artifacts can introduce DNA assembly errors in sequencing workflows (20). Thus, the 96 *T. kodakarensis* amplified fosmids were treated with T7 endonuclease I to remove branched DNA structures formed by the polymerase and prepared for multiplexed ONT sequencing using 96 barcodes as described in the Materials and Methods. The library was sequenced for 48 h on a GridION sequencer. Over 2.7 million reads were generated with an estimated N50 of 11.5 kb. The average and median number of reads per barcode was 21,414 and 18,944, respectively (Supplementary table 1).

Rolling circle amplification of circular templates with phi29 can generate sequence concatemers and chimeras that present challenges for DNA sequence assembly from long-read data (18). To address these issues, we developed a novel informatics pipeline termed ‘phiXXer’ (Figure 3). This pipeline tackles potential concatemers and chimeric reads by first trimming away regions that match the fosmid backbone before the reads are used as an input for *de novo* assembly (Supplementary figure 2). After assembly, the contig corresponding to the insert is again inspected and any remaining vector matches are trimmed, producing a linear sequence of cloned fosmid insert beginning at the MCS of the vector (Supplementary figure 3). Using this pipeline, a contig sequence for the cloned insert was successfully assembled for 89 of the 96 fosmids sequenced. Contig size ranged from 1,338 to 43,320 bp with a median size of 34,151 bp (Figure 4.A.). The depth of coverage for the assembled contigs ranged from 28X to 5,683X with a median value of 726X.

**Figure 3.**
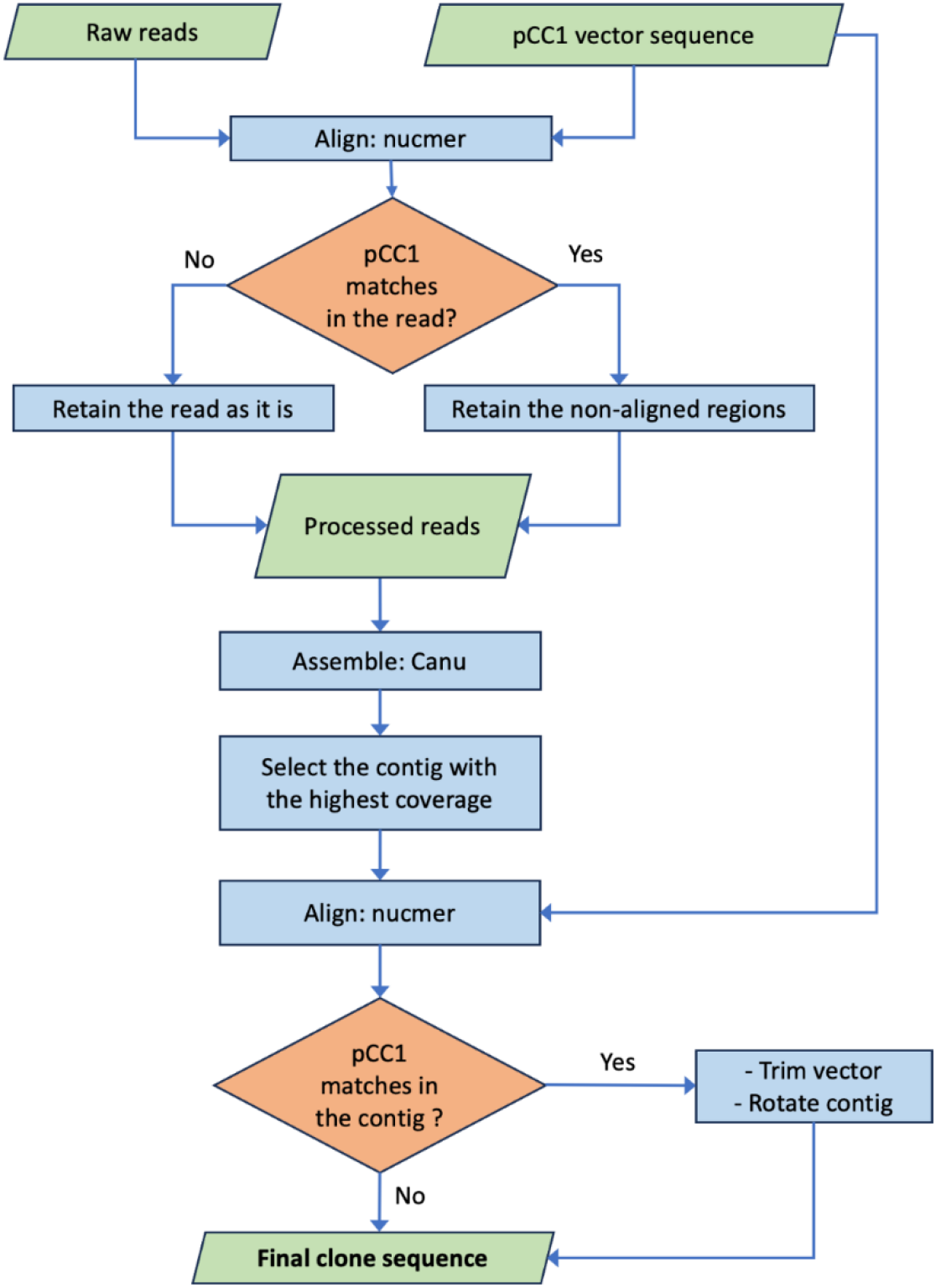
The phiXXer pipeline for *de novo* assembly of nanopore sequence data for phi29-amplified DNA. The common fosmid vector pCC1 was used in the illustrated protocol. Raw sequencing reads are aligned to the vector sequence and any matching regions are trimmed out before assembly. If multiple contigs are produced during the assembly, only the contig with the highest coverage is retained. This contig is aligned to the vector and any matching regions are trimmed away, producing a linear sequence of the cloned environmental or genomic DNA insert (free of the vector backbone). Nucmer is whole genome sequence aligner (21); canu is a long-read *de novo* assembler (22).

**Figure 4.**
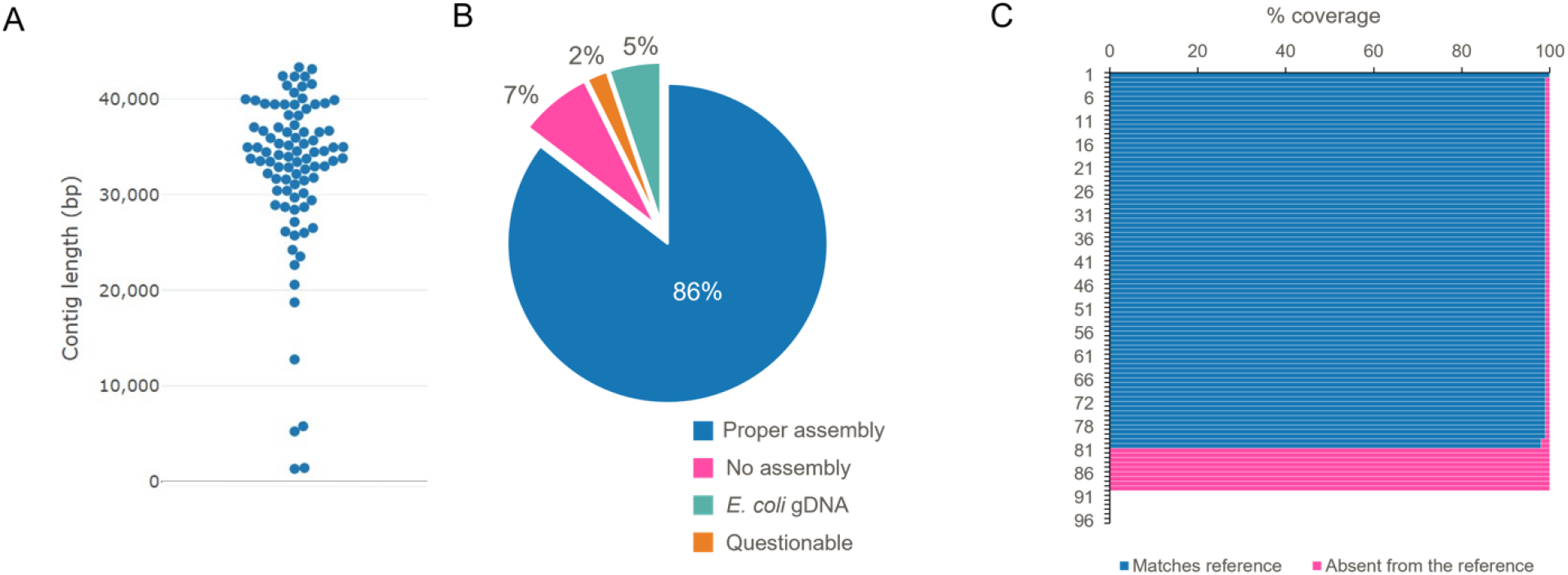
Assembly of 96 sequences from a *T. kodakarensis* genomic fosmid library using phiXXer. (A) Size distribution of the obtained contigs. (B) Evaluation of the assembly quality by comparing phiXXer contigs to reference sequences using BLASTN. ‘Proper assembly’: >99% coverage and 100% identity; ‘No assembly’: no contig generated; ‘*E. coli* gDNA’: contig matched *E. coli* genomic DNA; ‘Questionable’: contig matches *T. kodakarensis* genome but not the reference sequence. (C) Coverage percentage to the reference sequences by BLASTN for each fosmid.

To assess the accuracy of these 89 assemblies, each contig was compared using BLASTN to its corresponding reference nucleotide sequence that had been previously generated with the Illumina/plexWell™ technology. For 80 of 89 assembled fosmids, phiXXer output perfectly matched the reference data (>99% coverage and 100% identity) (Figure 4.A.B.C). In five instances, the generated contig did not match the reference fosmid but instead mapped to the *E. coli* genome. These contigs were significantly shorter (*e*.*g*., 1.3-12.8 kb with four contigs being below 6 kb) than those that matched a fosmid reference sequence (Figure 4.A.). Their presence in our sequencing library was likely caused by unspecific *E. coli* gDNA amplification or host gDNA carry-over from crude lysates. Finally, four phiXXer-generated contigs did not match their fosmid reference sequences but perfectly matched 30-40 kb loci elsewhere in the *T. kodakarensis* genome. Sanger sequencing the inserts from these fosmid showed that two of these contigs were correctly assembled by our workflow and that their Illumina reference sequences were incorrect. For the remaining two contigs, Sanger sequencing yielded a double sequence pattern, suggesting the fosmid backbone-specific primers had multiple priming sites. We hypothesize that these fosmids either contained two pCC1-FOS backbones (the result of a cos site bypass where cloned inserts are too short) or that two different fosmids were present in the same cell (Supplementary figure 4) (23).

Reproducibility of the phiXXer assembly pipeline was also evaluated. The nanopore sequencing reads generated for the 96 *T. kodakarensis* fosmids were independently assembled a total of five times and the results compared. In this evaluation, 82% of the fosmids were systematically assembled (*i*.*e*., a contig was generated in all five runs) while 95% yielded an assembled contig in four out of the five runs. The contig lengths for each assembled fosmid were also compared across each of the five phiXXer runs (Supplementary table 2). They were found to be identical (Δlength < 40 bp for a ∼35 kb contig) across all five runs for 77% of assembled fosmids. For 85% of the assemblies, contig lengths were found identical in at least four runs, and were identical in 92% of assemblies in at least three runs (Figure 5). Notably, out of the seven fosmids that did not yield assembly in our first run, five were successfully assembled and matched our reference in at least one of the other runs. Similarly, of the five contigs matching *E. coli* genome, four could be assembled in another run of phiXXer. A second phiXXer assembly run is therefore recommended for fosmids that do not assemble or yield a short contig during a first attempt.

**Figure 5.**
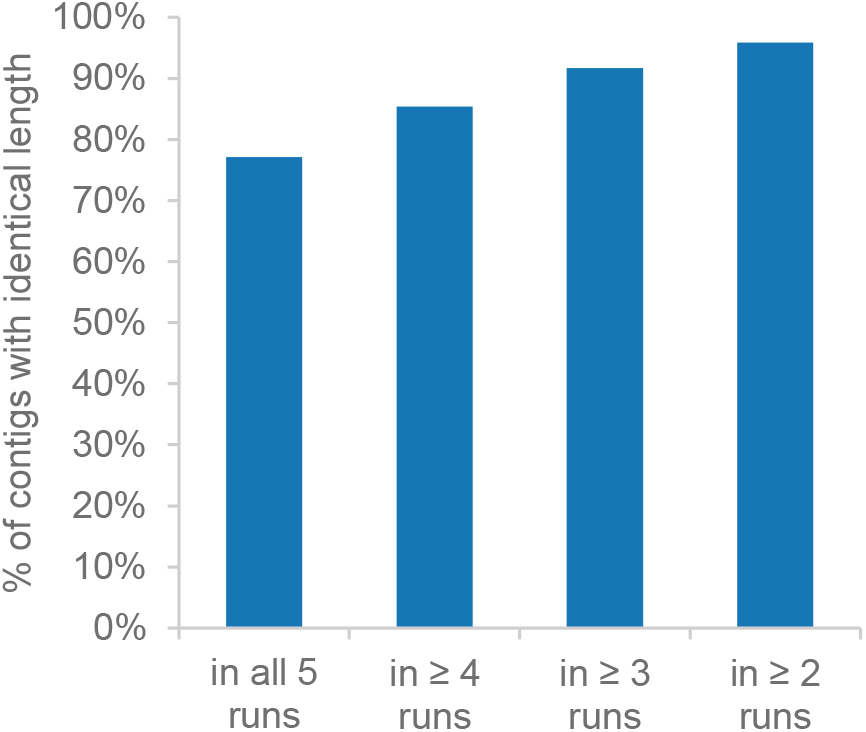
Percentage of identical contig lengths (Δlength<40 bp) across several runs of phiXXer. The pipeline phiXXer was run 5 times on demultiplexed reads from 96 *T. kodakarensis* genomic DNA fosmids (Supplementary table 2).

Considered together, our data support the conclusion that our workflow consisting of colony-based fosmid amplification, nanopore sequencing, and phiXXer sequence assembly was highly robust. Using a previously described genomic fosmid library, this workflow yields a ∼93% assembly rate. Of these, 92% perfectly matched their corresponding reference sequences generated using an orthogonal method.

### 3. Sequencing complex metagenomic fosmid clone libraries

To validate our method and pipeline on a more complex metagenomic fosmid library, we applied it to sequencing clones from a human gut microbiome (HGM) fosmid library that has previously been used in enzyme discovery screens (Chuzel et al., 2021; Fossa et al., 2023). A total of 1,436 fosmids were sequenced in 15 runs, with each run comprising 94-96 pooled fosmids. A contig was obtained for 1,331 fosmids corresponding to an assembly rate of ∼93%, consistent to that observed for assembly of *T. kodakarensis* genomic fosmids. Contig sizes ranged from 1.1 to 70.5 kb with a median size of 32.3 kb, the expected size of a typical fosmid insert (Figure 6). Only six contigs were larger than 50 kb showing phiXXer’s ability to de-concatenate reads, trim the vector and provide the insert sequence as final assemblies. The depth of coverage of the assembled contigs ranged from 7X to 24,000X, with a median value of 310X.

**Figure 6.**
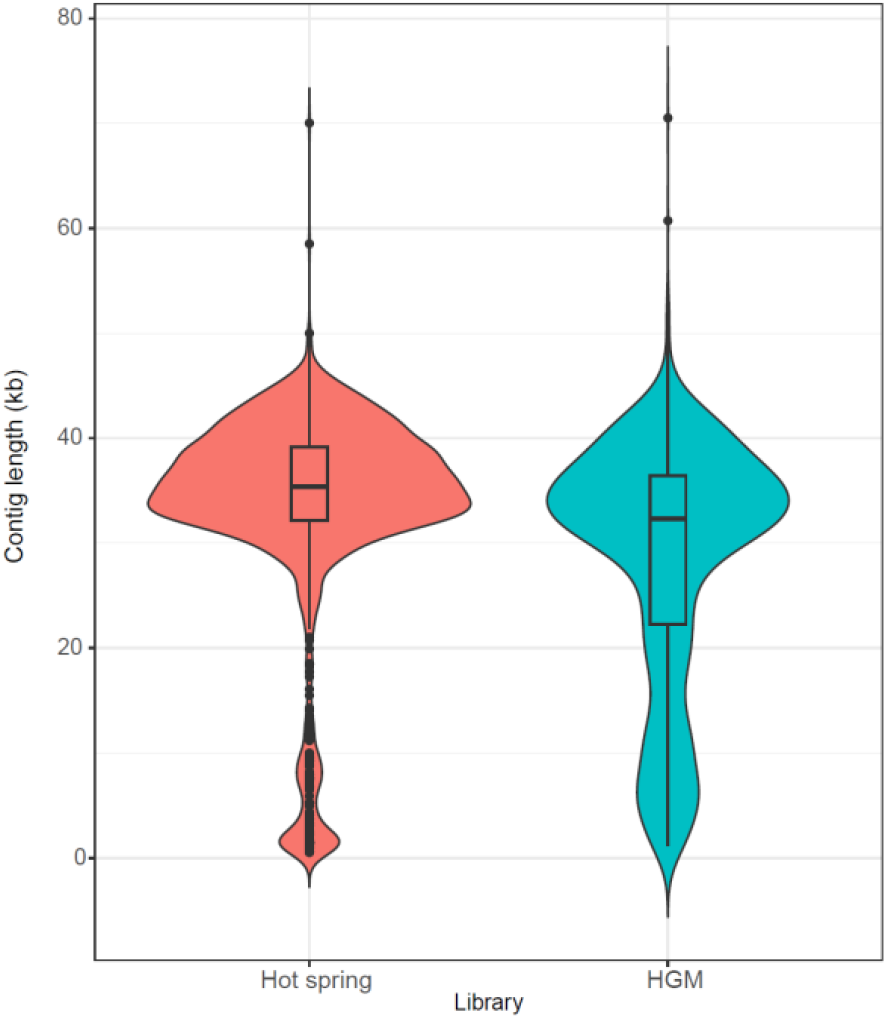
Contig size distribution using Oxford Nanopore Technologies/phiXXer compared to Illumina/seqWell. A total of 1,440 fosmids from a hot spring library were sequenced using Illumina and seqWell technology for barcoding and assembly. A total of 1,436 fosmids from a human gut microbiome library (‘HGM’) were sequenced using Oxford Nanopore Technologies and assembled using phiXXer. Size distribution of obtained contigs in both groups is illustrated using violin plots.

**Figure 7.**
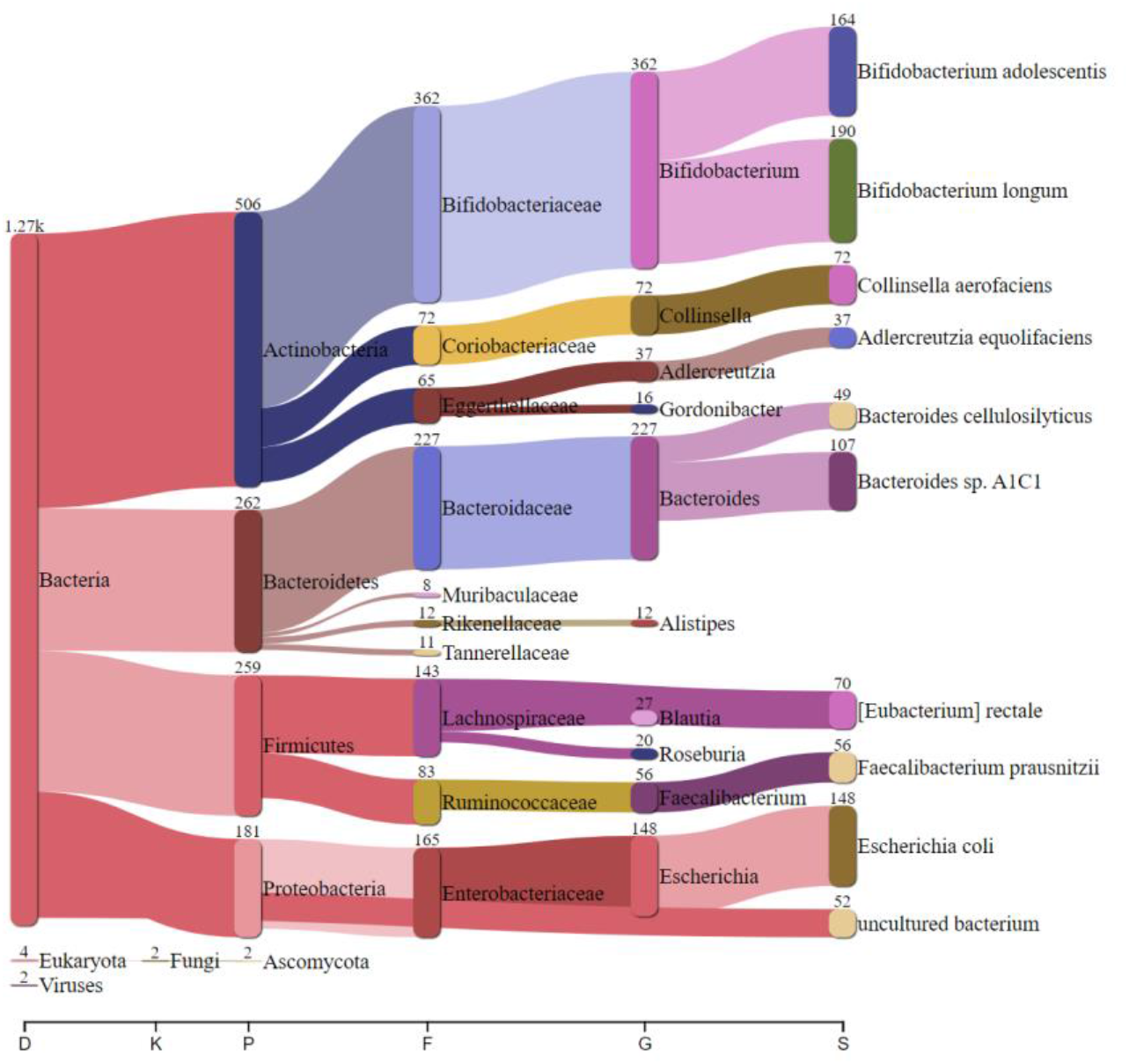
Taxonomic distribution of sequences from the human gut microbiome assembled fosmids. Metagenomic sequence classification with Kraken2 of 1,331 fosmids from our human gut microbiome library sequenced using Oxford Nanopore Technologies and assembled with phiXXer.

We also compared the insert size distribution of this complex HGM fosmid sequences obtained using our workflow to that of an independent set of complex metagenomic library of 1,440 fosmids generated from a hot spring microbiome and assembled using the Illumina/plexWell™ workflow. For this hot spring library, an assembly was generated for 1,384 out of 1,440 fosmids (96% assembly rate) with contigs ranging from 0.5 to 70.0 kb and a median of 35.4 kb (Figure 6). Even though the two fosmid libraries being compared are from different origins, their fosmid insert size distributions show a similar median and number of upper outliers. However, a larger number of contigs smaller than 20 kb are observed for the HGM library, an indication of potential *E. coli* gDNA contamination in our prepped ONT library.

To further evaluate the assembly quality of these previously un-sequenced HGM fosmids, a taxonomic analysis was performed using Kraken2 metagenomic sequence classifier (25). The vast majority of contigs were assigned to the domain Bacteria, consistent with the origin of the metagenomic library (26, 27). The most represented phyla were from Actinobacteria, Bacteroidetes, Firmicutes and Proteobacteria, in line with species found in the human gut microbiome of healthy individuals (26, 27). The contigs originating from *E. coli* could be derived from either the gut microbiome or a contaminant from the host strain gDNA. Upon further inspection, potential contaminant contigs were found to be less than 11% of all assembled contigs and exhibited shorter lengths and lower coverages (Supplementary figure 5). The contig classification is overall consistent with the human gut microbiome, suggesting proper fosmid sequencing and assembly.

## Discussion

In this work we present a novel approach for high-throughput fosmid sequencing that exploits phi29 polymerase to amplify fosmids directly from lysed bacterial cells and to generate material to prepare a sequencing library. Fosmids are sequenced using ONT making the workflow cost-effective and available to numerous labs. Up to 96 fosmids are sequenced in a single run ensuring a good throughput. We optimized the sample preparation to be straightforward eliminating the need for bead purification until fosmids are pooled together in a single tube. We also developed a bioinformatic pipeline termed phiXXer that enables easy and fast assembly of the reads to generate linearized fosmid insert sequences. phiXXer was developed to tackle some challenges inherent to both *de novo* assembly of circular molecules and phi29 products. The devised workflow was validated on more than a thousand fosmids from different sources.

The construction of fosmid metagenomic libraries and their screening has enabled the discovery of many enzymes and bioactive compounds (28–31). In these workflows, the ability to rapidly sequence fosmids from hits is imperative. We believe that the proposed approach to fosmid sequencing could speed up discoveries by providing a fast high-throughput sequencing and assembly method. With the present method enabling 96 fosmids to be sequenced and assembled from a single ONT run, larger screens (*e*.*g*., phenotypic screens) from more complex libraries could be considered where several thousand clones are surveyed, and hundreds of hits could be found. In addition, arrayed fosmid library collections could be pre-sequenced to create fosmid sequence databases, which would allow the retrieval of fosmid sequences corresponding to the hits immediately by using the clones’ coordinates. This would tremendously lower the ‘’hit to discovery timeframe’’. The proposed method could be expanded to plasmids, cosmids or more challenging vectors such as BACs and be applied to projects outside functional screenings where sequencing of these molecules is needed (32, 33).

Interestingly, we noticed that in some instances, multiple rounds of phiXXer are needed to obtain a proper fosmid assembly. Because *de novo* assemblers typically only use a subset of the generated data for initial seeding of the assembly, it is common to observe different contigs upon *de novo* assembly of the same dataset several times (34). The circular nature of the fosmid molecule and the known chimera formation artefacts from phi29 polymerase amplification could also contribute to non-deterministic assemblies (20). In future versions of phiXXer, one assembly iteration that yields a contig deemed of poor quality (low sequence coverage and/or short contig length) could inform a second iteration to improve assembly quality. Finally, the observed depth of coverage greater than 300X suggests that pooling more than 96 fosmids together in a single ONT run might be possible and could be evaluated. To that end, custom barcodes would have to be synthesized since 96 is the maximum number of barcodes currently offered by ONT in their Native Barcoding Kit.

## Materials and Method

### Material

Phi29-XT and phi29-XT Reaction Buffer (from E1603: phi29-XT RCA Kit), dNTP solution mix, SacI-HF, XhoI, rCutSmart™ buffer, T7 endonuclease I, NEBuffer™ 2, NEBNext® FFPE DNA Repair Mix, NEBNext® FFPE DNA Repair Buffer, NEBNext® Ultra™ II End Prep Enzyme Mix, Next® Ultra™ II End Prep Reaction Buffer, Blunt/TA Ligase Master Mix, NEBNext® Quick ligation reaction buffer, Quick T4 DNA ligase were all from New England Biolabs (Ipswich, MA).

UltraPure™ Bovine serum albumin (BSA) was from Invitrogen (Waltham, MA). Genomic DNA ScreenTape and reagents were from Agilent (Santa Clara, CA). Qubit™ BR dsDNA kit was from Thermo Fischer Scientific (Waltham, MA). CopyControl™ inducing solution was from LGC Biosearch Technologies (Hoddesdon, UK).

Primers were ordered from Integrated DNA Technologies (Coralville, IA) with five phosphorothioate bonds between the last 6 bases to prevent degradation by phi29-XT.

pCC1 forward: 5’-GGATGTGCTGCAAGGCGATTAA^*^G^*^T^*^T^*^G^*^G-3’

pCC1 reverse: 5’-CTCGTATGTTGTGTGGAATTG^*^T^*^G^*^A^*^G^*^C-3’

(where ^*^ denotes a phosphorothioate bond)

#### 1. Fosmid rolling circle amplification using phi29-XT

Ninety-six clones were spotted from glycerol liquid stocks onto a LB-agar plate containing 12.5 µg/mL chloramphenicol and 1X CopyControl™ inducing solution using a pin-replicator. Clones were grown overnight at 37°C to form arrayed colonies. Each colony was then scraped and resuspended in the corresponding well of a 96-well PCR plate containing 5 μL of 60 mM KOH, 2 mM EDTA pH 8.0 per well, using a multichannel pipette. Cells were lysed at 60°C for 10 min. Lysates were then neutralized by addition of 5 μL of 75 mM Tris-HCl, pH 7 per well. A master mix for fosmid amplification was prepared by mixing 440 μL of phi29-XT Reaction Buffer (5X), 220 μL of 10 mM dNTP solution mix, 110 μL of 8 μM pCC1 forward primer, 110 μL of 8 μM pCC1 reverse primer, 110 μL of phi29-XT (10X) and 110 μL of miliQ water. Each neutralized cell lysate was mixed with 10 μL of master mix and incubated at 42°C for 2-8 h followed by 65°C for 10 min in a thermocycler (lid at 75°C).

Amplified products were analyzed by agarose gel electrophoresis as follows. In a 25 μL total reaction volume, 2 μL of amplification reaction was mixed with 20 U each of SacI-HF and XhoI and 2.5 μL 10X rCutSmart™ buffer. Reactions were incubated for 1 h at 37°C followed by 10 min at 65°C. Digests were separated using a 0.8% agarose gel.

#### 2. ONT library preparation and sequencing

Amplified fosmids were first debranched using T7 endonuclease I to remove the branching DNA structures that are often created by phi29 (20). In a new 96-well PCR plate, 12 μL of each phi29-XT-amplified product were mixed with 1 μL of T7 endonuclease I, 2 μL of NEBuffer™ 2 and 5 μL of water. Reactions were incubated at 37°C for 2h.

Debranched samples were then treated to repair DNA and end damage. In a new 96-well PCR plate, 12 μL of each debranched sample was aliquoted. A repair master mix was prepared by combining 94.5 μL of NEBNext® FFPE DNA Repair Buffer, 94.5 μL of NEBNext® Ultra™ II End Prep Reaction Buffer, 81 μL NEBNext® Ultra™ II End Prep Enzyme Mix and 54 μL NEBNext® FFPE DNA Repair Mix. Three microliters of repair master mix were added to each well. Reactions were incubated in a thermocycler at 20°C for 5 min followed by 65°C for 5 min.

Repaired samples were then barcoded for multiplexing. In a new 96-well PCR plate, 1.5 μL of each end-repaired DNA sample was aliquoted and mixed with 2.25 µL of water, 1.25 µL of Native Barcodes (NB01-96) (Native Barcoding Kit 96 V14, Oxford Nanopore Technologies, UK) and 5 µL Blunt/TA Ligase

Master Mix for a total of 10 µL per reaction/well. Ligation was performed at room temperature for 40 min. Reactions were stopped by addition of 1 µL 0.5 M EDTA. All 96 samples were then pooled together in a single 1.5 mL microcentrifuge tube. The barcoded pool was then cleaned up with 384 µL of AMPure XP beads (0.4X volume ratio) (Native Barcoding Kit 96 V14, Oxford Nanopore Technologies, UK). DNA was allowed to bind to the beads for 10 min at room temperature on a rotator. The sample was then placed on a magnetic rack and the supernatant removed to isolate the beads. The beads were washed twice with 1 mL of 70 % freshly prepared ethanol. After the second wash, excess ethanol was removed carefully, and the sample air-dried for 30 s. The beads were then carefully resuspended in 35 µL of MiliQ water and incubated at 37°C for 10 min to elute the bound DNA. To improve elution, tube was flicked every 2 min.

Sequencing adaptors were then added by mixing 30 µL of the cleaned DNA pool with 5 µL of native adapter (Native Barcoding Kit 96 V14, Oxford Nanopore Technologies, UK), 10 µL of NEBNext® Quick ligation reaction buffer (5X) and 5 µL of Quick T4 DNA ligase. Ligation was performed at room temperature for 20 min. The sequencing library was purified using 20 µL AMPure XP beads (0.4X volume ratio). Binding was performed at room temperature for 10 min on a rotator. The sample was then placed on a magnetic rack and the supernatant removed to isolate the beads. The beads were washed twice with 125 µL of Long Fragment Buffer (Native Barcoding Kit 96 V14, Oxford Nanopore Technologies, UK). DNA was eluted with 15 µL of Elution Buffer (Native Barcoding Kit 96 V14, Oxford Nanopore Technologies, UK) at 37°C for 10 min. The concentration of the prepared sequencing library was then assessed using the Qubit™ BR dsDNA kit. The described protocol typically generates large fragment libraries (40-60 kb) and the DNA concentration of the final library was diluted to 60 ng/µL if needed. It is recommended to load 10-20 fmol of library on the nanopore flow cell.

The multiplexed library was run on a GridION using flow cells with the R10.4.1 chemistry following the manufacturer’s instructions. Briefly, the priming mix was prepared by mixing 5 µL of BSA 50 mg/mL with 30 µL of Flow Cell Tether and 1,170 µL of Flow Cell Flush (Native Barcoding Kit 96 V14, Oxford Nanopore Technologies, UK). Air was removed from the priming port and 800 µL of priming mix loaded into the priming port with the sequencing hole closed. After 5 min, an additional 200 µL of priming mix was loaded into the priming port with the sequencing hole opened this time. Finally, the sequencing mix comprised of 12 µL of multiplexed library, 25.5 µL of Library Beads and 37.5 µL of Sequencing Buffer (Native Barcoding Kit 96 V14, Oxford Nanopore Technologies, UK) was loaded drop by drop into the sequencing hole. Both the priming port and sequencing hole were closed and the run performed with the following settings: barcode trimming, super-accuracy basecalling, 260 bps (accuracy mode) for 48-72h.

#### 3. The phiXXer pipeline for amplified fosmid insert assembly

De-multiplexed raw reads from the GridION sequencer were obtained using MinKNOW. For the phiXXer pipeline, the pCC1 vector reference sequence used in the vector trimming steps was obtained by rotating the sequence such that the multiple-cloning site (MCS) was at the 3’ prime end of the sequence. For each clone, the reads were individually aligned to the rotated pCC1 sequence using nucmer version 3.1. (21). If a pCC1 match was found in the read, the aligned region was trimmed out, while the non-matching flanking regions were retained as separate final reads (Supplementary figure 2). The reads without pCC1 alignments were retained as such. Any reads or read-fragments shorter than 150 nucleotides were discarded. The processed reads were assembled using canu version 2.2 (22) with default parameters, except the parameters “stopOnLowCoverage” and “minInputCoverage” were set to a value of 7. The canu contig with the highest length and coverage was selected as the insert sequence for each clone. This contig was inspected for any alignments to the pCC1 backbone and any vector matching region was trimmed out as described above, while the flanking regions were merged into a consensus such that they are consistent with the expected 5’ to 3’ orientation in the MCS (Supplementary figure 3). Quality metrics such as the mean depth of coverage and percent of sequenced reads contributing to the assembled contig were obtained by mapping the reads to the insert sequence using minimap2 version 2.24-r1122 (35) and analysis of the bam files using samtools version 1.18 (36). Taxonomic classification of the insert sequences obtained from the phiXXer pipeline was performed using kraken version 2.0.9-beta (37) based on the NCBI nt database. To search for contigs potentially originating from the *E. coli* host, a megablast search was performed against the reference genome of strain NEB10-beta (GenBank accession CP053604.1). The contigs with greater than 95% sequence identity over 90% of their length were marked as potentially arising from host gDNA contamination.

## Acknowledgements

We thank Dr. Donald Comb and New England Biolabs for basic research support. We also thank Samantha Fossa, Dr. Dora Posfai, Dr. Kyle Vrtis, for comments and suggestions on the manuscript, Dr. Yanxia Bei for technical expertise, Dr. Vladimir Potapov, Dr. Zhiyi Sun and Dr. Sean R. Johnson for helpful discussions and Dr. Philipp C. Schneider for supports with RStudio.

