## Supplementary information for "High-throughput nanopore DNA sequencing of large insert fosmid clones directly from bacterial colonies"

### Supplementary Tables

| *Thermococcus kodakarensis* fosmids | | | |
|---|---|---|---|
| **barcode** | **# reads** | **barcode** | **# reads** |
| barcode01 | 19019 | barcode50 | 23163 |
| barcode02 | 74918 | barcode51 | 44257 |
| barcode03 | 39761 | barcode52 | 12021 |
| barcode04 | 24825 | barcode53 | 33380 |
| barcode05 | 10947 | barcode54 | 34240 |
| barcode06 | 14385 | barcode55 | 15353 |
| barcode07 | 3000 | barcode56 | 22985 |
| barcode08 | 15076 | barcode57 | 20266 |
| barcode09 | 8602 | barcode58 | 46382 |
| barcode10 | 42070 | barcode59 | 31816 |
| barcode11 | 35009 | barcode60 | 19058 |
| barcode12 | 27993 | barcode61 | 18827 |
| barcode13 | 21820 | barcode62 | 38807 |
| barcode14 | 10519 | barcode63 | 10329 |
| barcode15 | 6883 | barcode64 | 17684 |
| barcode16 | 15200 | barcode65 | 27791 |
| barcode17 | 5992 | barcode66 | 5273 |
| barcode18 | 18355 | barcode67 | 10072 |
| barcode19 | 39015 | barcode68 | 13871 |
| barcode20 | 4795 | barcode69 | 25762 |
| barcode21 | 12529 | barcode70 | 19827 |
| barcode22 | 16413 | barcode71 | 8363 |
| barcode23 | 3015 | barcode72 | 19253 |
| barcode24 | 32522 | barcode73 | 10406 |
| barcode25 | 26003 | barcode74 | 42447 |
| barcode26 | 30454 | barcode75 | 31959 |
| barcode27 | 40445 | barcode76 | 10652 |
| barcode28 | 40480 | barcode77 | 10214 |
| barcode29 | 24322 | barcode78 | 22899 |
| barcode30 | 20221 | barcode79 | 10430 |
| barcode31 | 29106 | barcode80 | 22741 |
| barcode32 | 24297 | barcode81 | 12440 |
| barcode33 | 14427 | barcode82 | 33998 |
| barcode34 | 12794 | barcode83 | 28747 |
| barcode35 | 33936 | barcode84 | 11338 |
| barcode36 | 15905 | barcode85 | 25316 |
| barcode37 | 21915 | barcode86 | 16038 |
| barcode38 | 26496 | barcode87 | 3733 |
| barcode39 | 4745 | barcode88 | 11577 |
| barcode40 | 8502 | barcode89 | 16731 |
| barcode41 | 23141 | barcode90 | 40898 |
| barcode42 | 35013 | barcode91 | 30272 |
| barcode43 | 31779 | barcode92 | 7201 |
| barcode44 | 6633 | barcode93 | 5471 |
| barcode45 | 18869 | barcode94 | 8448 |
| barcode46 | 8095 | barcode95 | 16385 |
| barcode47 | 4814 | barcode96 | 79830 |
| barcode48 | 11636 | unclassified | 218282 |
| barcode49 | 10120 | | |
| average # reads/barcode | | | 21414.19 |
| Median # reads/barcode | | | 18944 |

*Supplementary table 1. Read distribution per barcode upon sequencing of 96 Thermococcus kodakarensis fosmids using Oxford Nanopore Technologies.*
*Reads were demultiplexed using MinKNOW.*

| Clone barcode | Contig length (bp) | | | | | Clone barcode | Contig length (bp) | | | | |
|---|---|---|---|---|---|---|---|---|---|---|---|
| | **run 1** | **run 2** | **run 3** | **run 4** | **run 5** | | **run 1** | **run 2** | **run 3** | **run 4** | **run 5** |
| 1 | 28714 | 28714 | 28714 | 28720 | 28714 | 49 | 33758 | 33757 | 33758 | 33758 | 33757 |
| 2 | 32655 | 32654 | 32655 | 32655 | 32655 | 50 | 42346 | 42375 | 42376 | 42376 | 33999 |
| 3 | 1431 | 41309 | 41309 | 41309 | 41309 | 51 | 18745 | 5926 | 18745 | 4937 | 1305 |
| 4 | 33527 | 33527 | 33528 | 33527 | 33527 | 52 | 33513 | 33513 | 33513 | 33512 | 33513 |
| 5 | 34557 | 34557 | 34558 | 34556 | 34558 | 53 | 40663 | 7187 | | 5577 | 4255 |
| 6 | 36534 | 36535 | 36535 | 3247 | | 54 | 41415 | 41413 | 41415 | 41417 | 41416 |
| 7 | 30417 | 30416 | 30416 | 30416 | 30416 | 55 | 31772 | 31772 | 31772 | 31772 | 31772 |
| 8 | 35316 | 35318 | 35317 | 35317 | 35317 | 56 | 37033 | 37033 | 37033 | 37033 | 37033 |
| 9 | 26145 | 26144 | 26144 | 26144 | 26144 | 57 | 37048 | 37048 | 37050 | 37048 | 37048 |
| 10 | 32969 | 32969 | 26916 | 32969 | 32969 | 58 | 1338 | 1326 | | 30729 | 1217 |
| 11 | 23540 | 23535 | 23544 | 23536 | 23547 | 59 | 38320 | 38320 | 38320 | 38320 | 7201 |
| 12 | 32947 | 32947 | 32947 | | 5881 | 60 | 36680 | | | 36680 | |
| 13 | 38283 | 38283 | 38283 | 38283 | 38283 | 61 | 42403 | 42405 | 5513 | 42404 | 42406 |
| 14 | 26014 | 26015 | 1341 | 3154 | 26010 | 62 | 34448 | 34448 | 34448 | 34448 | 34448 |
| 15 | 24226 | 24226 | 24205 | 24226 | | 63 | 30155 | 30155 | 30155 | 30155 | 30155 |
| 16 | 39447 | 39447 | | 39447 | 39447 | 64 | 39975 | 39975 | 39975 | 39975 | 39975 |
| 17 | 28905 | 28905 | 28471 | 28905 | 28905 | 65 | 43320 | 43320 | 43320 | 43320 | 43320 |
| 18 | 34447 | 34447 | 34447 | 34447 | 34447 | 66 | 41560 | 41592 | 41593 | 41592 | 41593 |
| 19 | 35942 | 35942 | 35942 | 35942 | 35942 | 67 | 35917 | 35917 | 35917 | 35917 | 35917 |
| 20 | | | 24786 | 24786 | 24786 | 68 | | 33834 | 1326 | 5628 | 5155 |
| 21 | | 17645 | 17644 | 1337 | 17645 | 69 | 33963 | 33963 | 33963 | 33963 | 33963 |
| 22 | 38963 | 38963 | 38962 | 38963 | 38963 | 70 | 32159 | 32159 | 32159 | 32159 | 32159 |
| 23 | 39893 | 39893 | 39893 | 39893 | 39893 | 71 | 31479 | 31479 | 31479 | 31479 | 31479 |
| 24 | 34952 | 34952 | 34952 | 34952 | 3315 | 72 | 41309 | 41309 | 41309 | 41309 | 41309 |
| 25 | 33806 | 33806 | 1206 | 33806 | 33806 | 73 | 31593 | 31593 | 31593 | 31593 | 31593 |
| 26 | 35150 | 35150 | 35150 | 35150 | 35150 | 74 | 31660 | 31659 | 31661 | 31661 | 31661 |
| 27 | 34934 | 34935 | 34935 | 34935 | 34935 | 75 | 39411 | 39410 | 39411 | 39423 | 39411 |
| 28 | 34981 | 34981 | 34981 | 34981 | 34981 | 76 | 34538 | 34538 | 34533 | 34526 | 34538 |
| 29 | | | 1198 | 1200 | | 77 | 36536 | 36547 | 36537 | 36537 | 36537 |
| 30 | 33446 | 33446 | 33446 | 33446 | 33446 | 78 | 30412 | 30413 | 30413 | 30413 | 30413 |
| 31 | 36664 | 36670 | 36668 | 36670 | 36655 | 79 | 29704 | 29704 | 29704 | 29704 | 29704 |
| 32 | 22647 | 22647 | 22647 | 22647 | 22647 | 80 | 39438 | | | 39440 | |
| 33 | 34151 | 34151 | 34151 | | 34151 | 81 | 31079 | 31079 | 31078 | 31079 | 31079 |
| 34 | 5793 | 21714 | 21714 | 21714 | | 82 | 43126 | 43125 | 43114 | 43126 | 43113 |
| 35 | 25735 | 25735 | 25735 | 25735 | 25735 | 83 | 37282 | 37282 | 37282 | 37282 | 37282 |
| 36 | 32851 | 32851 | 32851 | 32851 | 32850 | 84 | | 31018 | 31020 | 31017 | 31017 |
| 37 | 35337 | 35337 | 35337 | 35337 | 35337 | 85 | 39507 | 39508 | 39507 | 39507 | 39507 |
| 38 | 28701 | 28701 | 28701 | 28701 | 28701 | 86 | 26516 | 26516 | 26528 | 26528 | 26515 |
| 39 | 27151 | 27151 | 27151 | 27151 | 27151 | 87 | 40067 | 40077 | 40067 | 40078 | 40080 |
| 40 | 39560 | 39560 | 39560 | 39560 | 39560 | 88 | 28426 | 28413 | 28425 | 28426 | 28426 |
| 41 | 39849 | 39850 | 39850 | 39850 | 39850 | 89 | | 3980 | | 39140 | 39142 |
| 42 | | 32864 | 5625 | 32252 | 32864 | 90 | 42355 | 42367 | 42367 | 42367 | 42367 |
| 43 | 36525 | 36520 | 36519 | 36520 | 36520 | 91 | 34941 | 34941 | 34954 | 34953 | 34954 |
| 44 | 33768 | 28998 | 33769 | 33769 | 33770 | 92 | 12774 | 13846 | 10016 | 10086 | 8149 |
| 45 | 32884 | 32884 | 32884 | | 32884 | 93 | 29412 | | 29400 | 29412 | 29399 |
| 46 | 20588 | 20588 | 20588 | 20588 | 20588 | 94 | 35665 | 35667 | 35664 | 35664 | 35654 |
| 47 | 32216 | 32216 | 32216 | 32216 | 32216 | 95 | 33418 | 33430 | 33418 | 33429 | 33430 |
| 48 | 5251 | 40698 | 4569 | 1325 | 40698 | 96 | 39429 | 39434 | 39434 | 39420 | 39433 |

*Supplementary table 2. phiXXer reproducibility.*
*The phiXXer pipeline was executed 5 times to assemble sequencing reads from 96 Thermococcus kodakarensis fosmids (one fosmid per barcode) generated using Oxford Nanopore Technologies. The length of generated contigs is reported for each assembly run and each barcode.*

### Supplementary Figures

A

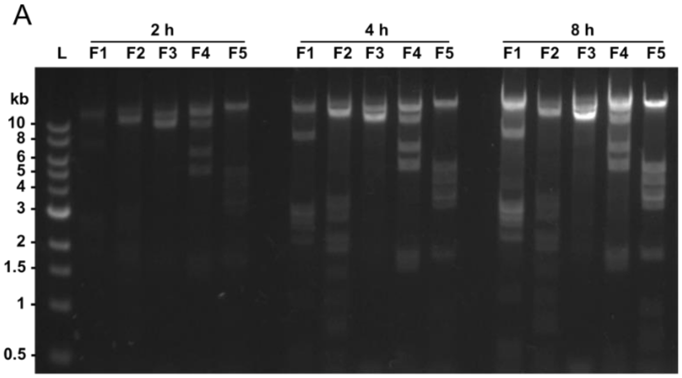


B

| ng/µL | 2 h | 4 h | 8 h |
|---|---|---|---|
| **F1** | 41.1 | 104 | 325 |
| **F2** | 57.7 | 134 | 158 |
| **F3** | 61.9 | 185 | 313 |
| **F4** | 106 | 226 | 332 |
| **F5** | 235 | 108 | 568 |

*Supplementary figure 1. Fosmid amplification using phi29 polymerase for different incubation times. Five different clones (F1-F5) from a T. kodakarensis genomic fosmid library were amplified for 2h, 4h or 8h using phi29-XT. (A) Amplification products were digested with SacI and XhoI and digested products analyzed by separation on a 0.8% agarose gel. (B) DNA concentrations obtained after amplification were measured with Qubit dsDNA assay.*

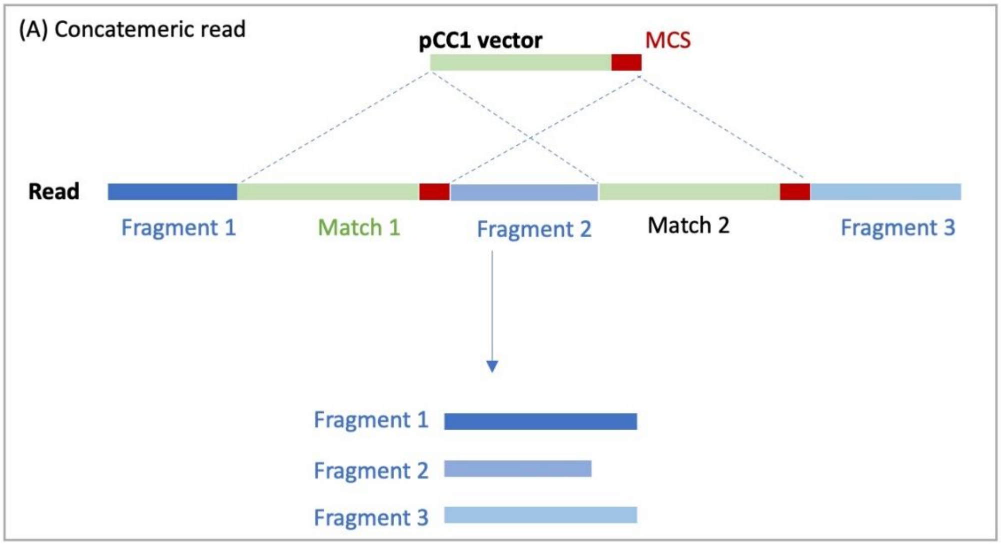


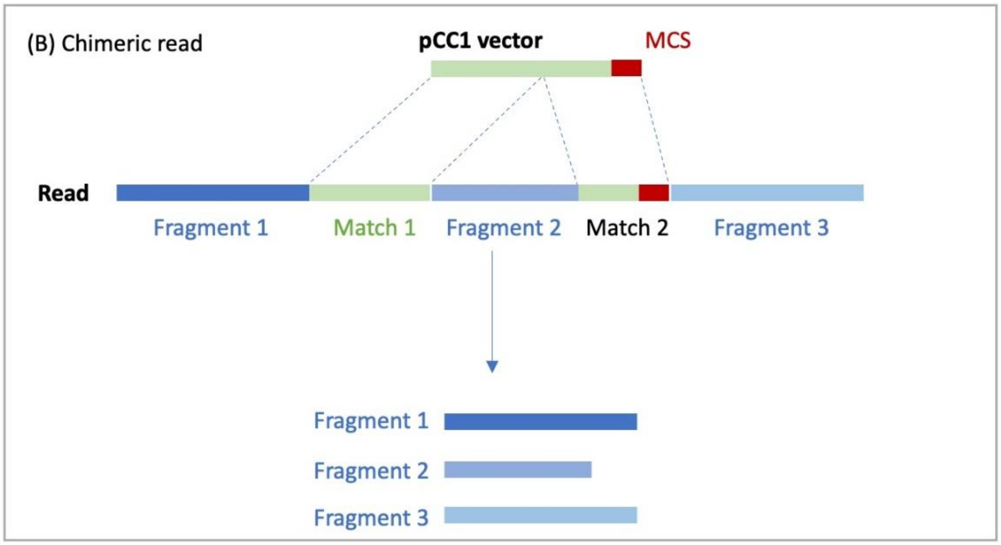


*Supplementary figure 2. Removing the vector sequences from the raw reads. The vector may be found as (A) multiple repeats with a concatemer read produced by rolling circle amplification, or (B) multiple matching segments due to chimera formation inherent in phi29 polymerase activity.*

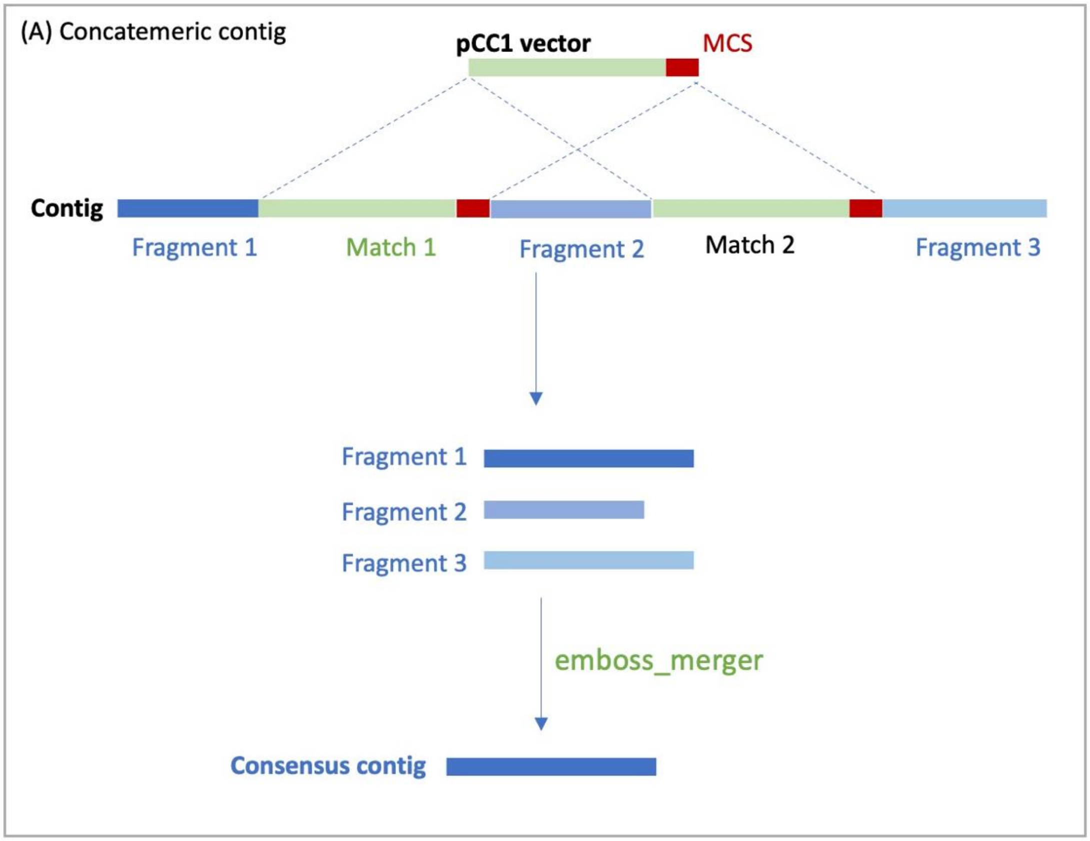


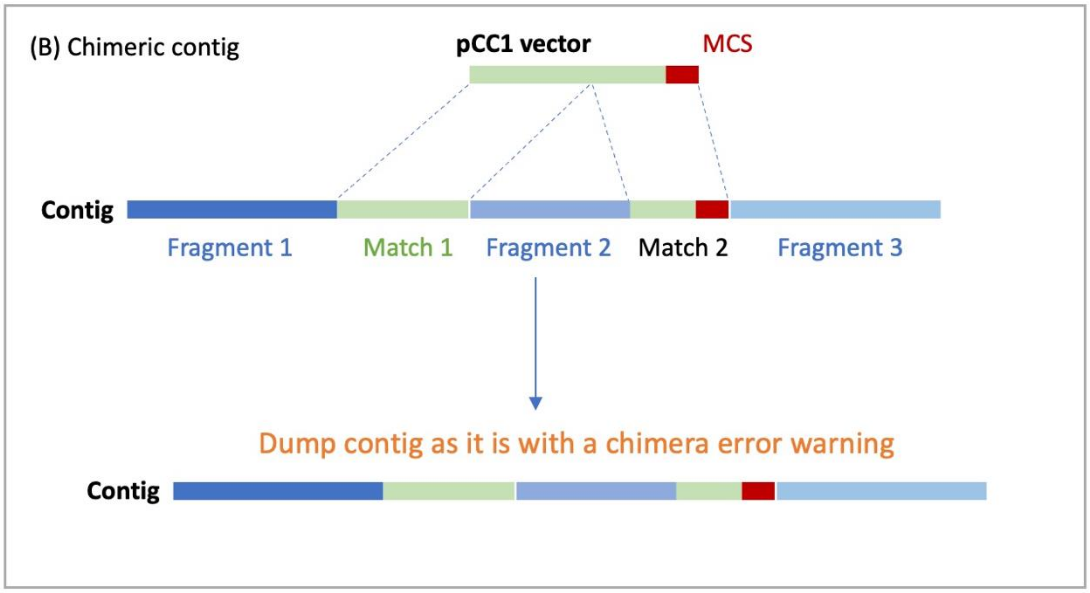


*Supplementary figure 3. Vector trimming from assembled contig (A) If a concatemeric contig is produced, the non-vector sequence regions can be merged into a single unique consensus sequence that begins at the MCS site. (B) In case of chimeric assembly, more than one complete or partial matches to the vector might be found on the assembled contig, and obtaining a single consensus sequence fails. The assembled contig is included in the output with an error message in fasta description to alert the user to a possible chimera.*

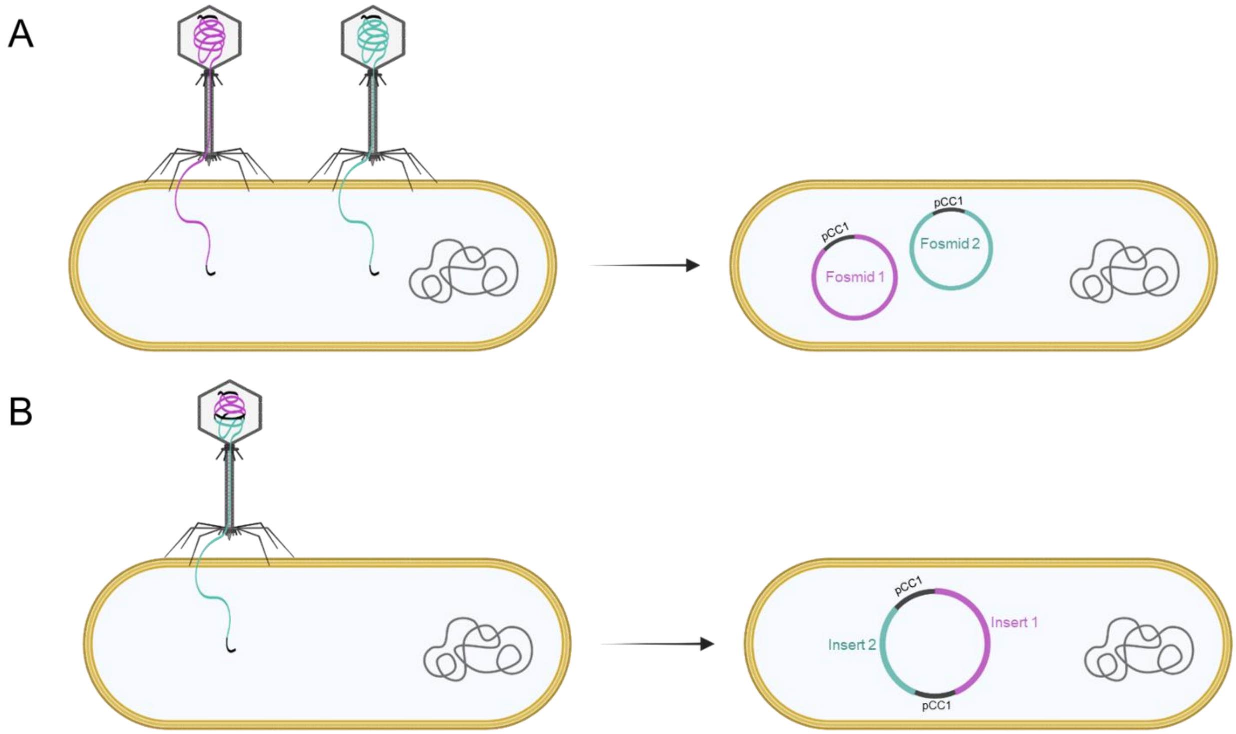


*Supplementary figure 4. Hypotheses on two distinct phage-mediated fosmid transduction mechanisms resulting in either two distinct fosmids in the same cell (A) or two fosmid backbones (pCC1) within a single fosmid (B). In (A) two unique lambda phage particles infect the same cell resulting in the insertion of two distinct fosmids in an Escherichia coli cell. In (B) short fosmid inserts (~15-20kb) are ligated to pCC1. The distance between two cos sites (located on pCC1) is too short to enable packaging in lambda phage particle resulting in the bypass of one cos site. Ultimately, two pCC1 backbones are packed with two distinct inserts. In vivo re-circularization forms a single circular fosmid. In both presented mechanisms, fosmids isolated from the cells cannot be end-sequenced using fosmid backbone (pCC1) specific primers as two binding sites are present.*

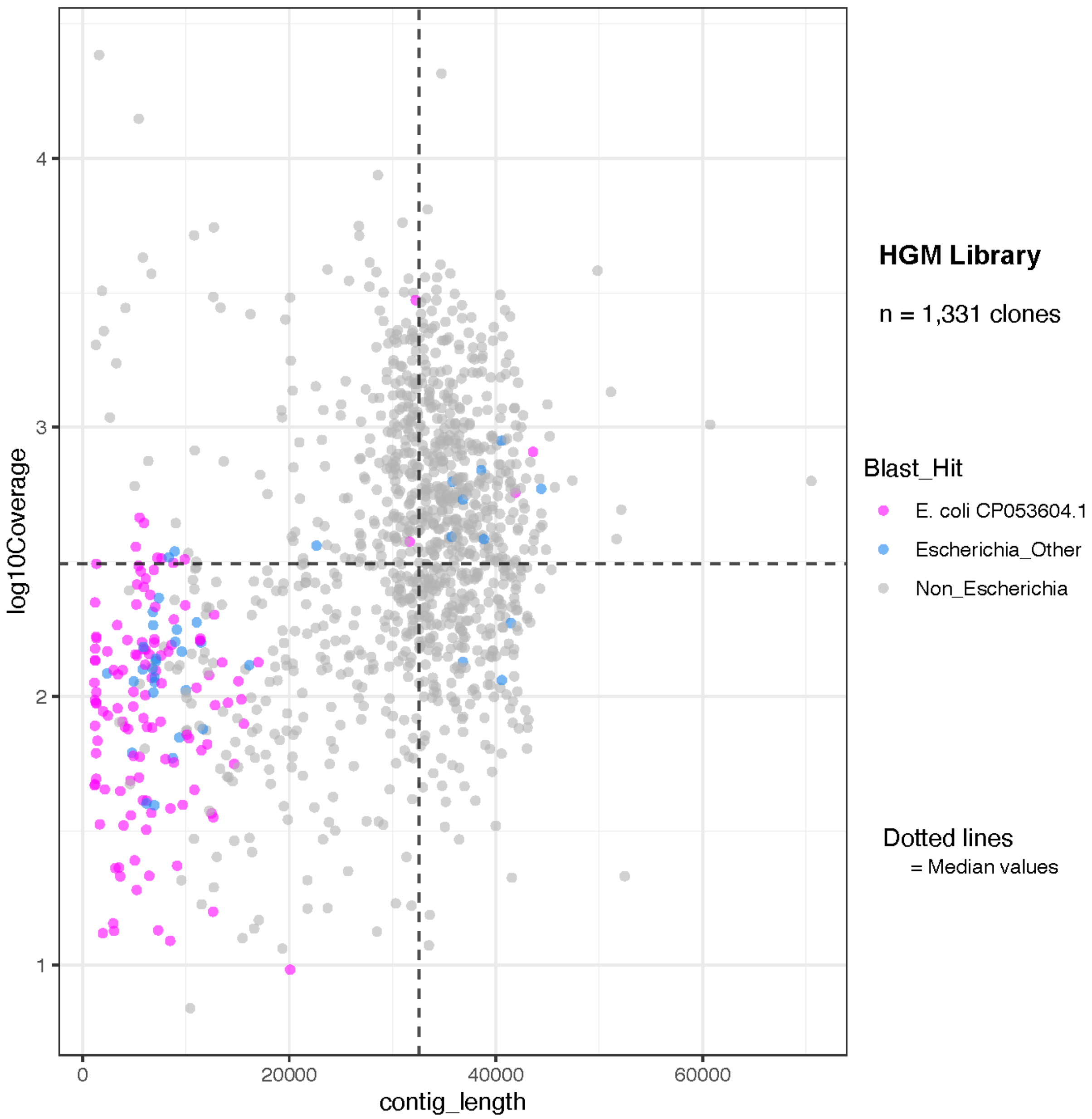


*Supplementary figure 5. Quality checks for the human Gut microbiome (HGM) library. Mean depth of coverage for each assembled insert contig was obtained from mapping the reads to the contig. Taxonomic classifications were obtained using kraken2. The contigs originating from the host E. coli strain NEB10-beta were identified using megablast against the reference genome accession CP053604.1 Most of the contigs from the E. Coli host were observed to have lengths as well as converge depths lower than the median values for the entire dataset.*
